# AF3Complex Yields Improved Structural Predictions of Protein Complexes

**DOI:** 10.1101/2025.02.27.640585

**Authors:** Jonathan Feldman, Jeffrey Skolnick

## Abstract

**Motivation:** Accurate structures of protein complexes are essential for understanding biological pathway function. A previous study showed how downstream modifications to AlphaFold 2 could yield AF2Complex, a model better suited for protein complexes. Here, we introduce AF3Complex, a model equipped with both similar and novel improvements, built on AlphaFold 3.

**Results:** Benchmarking AF3Complex and AlphaFold 3 on a large dataset of protein complexes, it was shown that AF3Complex outperforms AlphaFold 3. Moreover, by evaluating the structures generated by AF3Complex on datasets of protein-peptide complexes and antibody-antigen complexes, it was established that AF3Complex could create high-fidelity structures for these challenging complex types. Additionally, when deployed to generate structural predictions for protein complexes used in the recent CASP16 competition, AF3Complex yielded structures that would have placed it among the top models in the competition.

**Availability:** The AF3Complex code is freely available at https://github.com/Jfeldman34/AF3Complex.git.

**Contact:** Please contact.

## Introduction

Underpinned by a near-complete corpus of structures for single-domain proteins along with the plethora of data in protein sequence databases, AlphaFold 2, a highly accurate deep learning-based approach to predicting protein structures from amino acid sequences unprecedented in its ability to model single and multi-domain proteins with high fidelity, reinvigorated and led to the rapid progress of protein-folding research [1, 2, 3]. Of the myriad research studies launched after the advent of AlphaFold 2, many aimed to improve upon the underlying processes and architecture of the model to enhance the accuracy of its structural predictions–advances that would prove to be immensely impactful in the model’s clinical applications, where precision is key [4, 5, 6]. Among those studies, one, in particular, developed a derivative of AlphaFold 2, called AF2Complex, which significantly improved upon the former model’s ability to predict the structure of protein-protein complexes, which are highly relevant biologically and clinically [4]. When released, AF2Complex provided state-of-the-art performance for protein-protein complex modeling, outperforming even AlphaFold-Multimer, a model specialized for that task [4, 7].

In May 2024, the newest version of the AlphaFold suite of models, AlphaFold 3, was released. The architecture of the AlphaFold 3 model deviated from the previous models of its kind tremendously. It employed a diffusion-based architecture that imbued it with the abilities of a generative artificial intelligence model and minimized the importance of the multiple sequence alignments (MSAs) within the model; relegating it to a peripheral role in the inference pipeline [8]. This new architecture has granted AlphaFold 3 several impressive new capabilities. AlphaFold 3, unlike AlphaFold 2 or AlphaFold-Multimer, can predict the structures of proteins with bound ligands and ions, including them in the final structural prediction, and those of protein-nucleic acid complexes, which are complexes formed by one or more proteins interacting with one or more molecules of DNA or RNA [2, 8]. Additionally, AlphaFold 3 outperforms the previous generation of AlphaFold models on all modeling tasks, including protein-protein complex structure prediction [8].

It remained unclear, however, as to how well AlphaFold 3 would perform when compared to specialized models from outside of its model suite, such as the aforementioned AF2Complex model. This question is especially vital to protein complex modeling, where minute differences in the structure, especially at the interface between protein chains, may lead to drastically different behaviors.

Thus, preliminarily, AF2Complex and AlphaFold 3 were compared on a dataset of 1091 protein complexes. This analysis showed that AlphaFold 3 was the clear victor in terms of accuracy, beating AF2Complex in both macroscopic structural accuracy, measured using the TM-Score [9] and lDDT [10] score, and in interfacial structural accuracy, evaluated using the DockQ score [11].

Since AlphaFold 3 was superior to AF2Complex and could model protein complexes with additional ligand or ion information, it seemed appropriate to test whether a derivative model could be developed from AlphaFold 3, as AF2Complex was built from AlphaFold 2. This model, AF3Complex, would apply the improvements of AF2Complex to the AlphaFold 3 model, taking advantage of the newer model’s advances in the accuracy of structural predictions and the scope of macromolecules it can model [8].

In this study, we examined to what extent the retrofitting of AlphaFold 3 with a modified MSA generation algorithm, one which wholly omits the paired MSA, and a specialized confidence score that examines the interfacial residues in a protein complex more accurately could improve the ability of the model to predict the structure of protein complexes. Those modifications were the very same ones made to AlphaFold 2 to develop AF2Complex [4]. However, this study also explores a third change: generating two sets of structural models, one with ligands or ions, if such information is included in the model input, and one without ligands or ions, to examine which model the deep learning framework is more confident in. Together, these three downstream modifications, when applied to the AlphaFold 3 model, yielded the AF3Complex model, which was capable of outperforming the former across several metrics of total and interfacial accuracy and modeling challenging protein complex structures. These included protein-peptide interactions and antibody-antigen interactions, which previous models struggled to properly predict [12].

## Methods

The design of the AF3Complex model pipeline is shown in Figure 1. Given an input protein or nucleic acid sequences with optional ligand and ion information, the AF3Complex pipeline generates only unpaired MSAs based on the MSA generation algorithm used by AlphaFold 3 [2, 8, 7]. The removal of the MSA pairing algorithm is more efficient as it omits a computationally costly procedure in the pipeline and allows the model to avoid the pitfalls of generating orthologous sequences across different species–a process that can be confounded by the presence of genetic paralogs, cross-talk between protein signaling pathways, and the assimilation of pathogenic genetic material, such as that from lysogenic viruses [4, 13]. Moreover, since paralogs are often precursors to the diversification of protein function through evolutionary processes, excluding the paired MSAs imbues the model with greater flexibility, as it extricates stringent and possibly inaccurate paired structures [4]. Once the MSAs and the structural templates are generated, they are passed into the AlphaFold 3 model backbone along with the protein and nucleic acid sequences and optional ligand and ion information. This information is embedded and processed, as is standard in AlphaFold 3, before being passed into the AlphaFold 3 Pairformer and diffusion subunits, which then output the possible protein complex structures, from which the best is chosen using the predicted Interface-Similarity (pIS) score, a score that, like the pTM or ipTM metric utilized by the AlphaFold 3 model, represents higher fidelity models at greater values, with a maximum value of 1 [8]. Lastly, if the data input into the pipeline includes ligands or ions, the model inference pipeline is run again. The model reuses the already generated feature information from the first iteration where ligand and ion information was included, but this time with this information omitted. This allows the model to generate a structural prediction with ligands and ions excluded in an expedient manner, as identical feature generation need be repeated. The outputs–those with ligands or ions and those without ligands or ions–are compared, and the model with the highest pIS is chosen. In other words, the AlphaFold 3 model backbone reprocesses the inputs with all ligands and ions excluded, thereby yielding models without any such information.

**Fig. 1.**
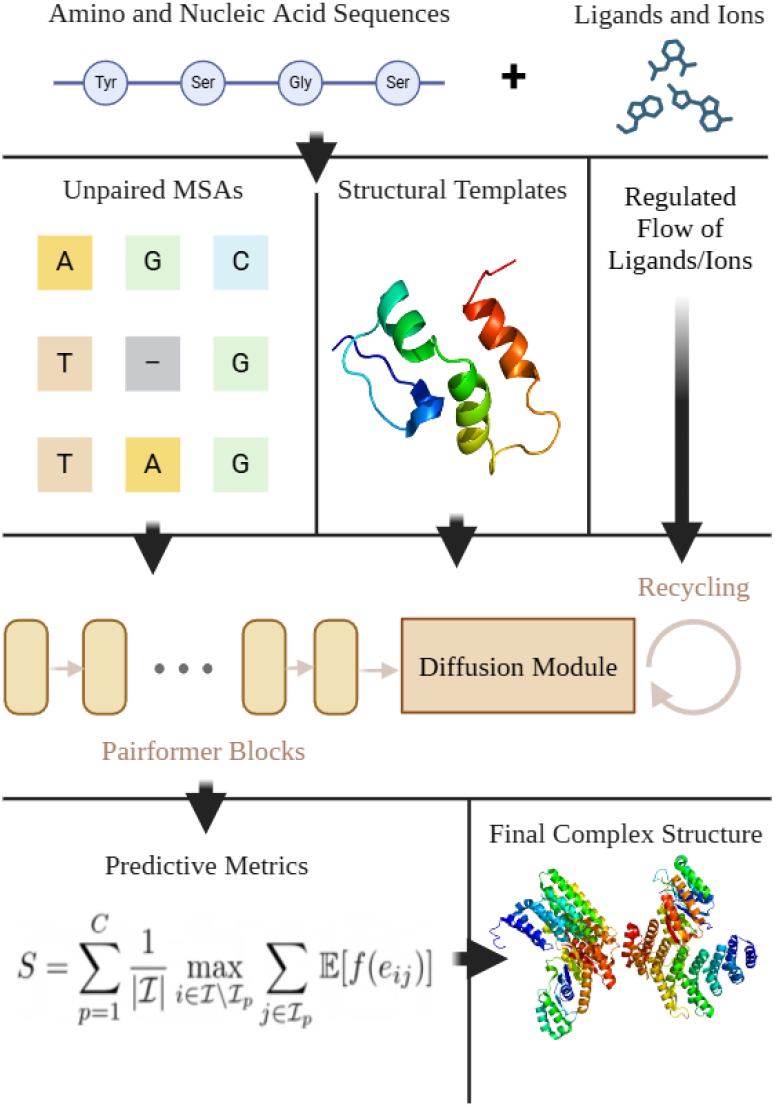
A schematic overview of the AF3Complex model pipeline, including the modified MSA generation algorithm, the AlphaFold 3 model backbone, the modified scoring metric, pIS, denoted S in the graph, and the regulated flow of ligand and ion information based on model confidence.

### AF3Complex Pipeline Overview

### Protein Complex Dataset Curation

The dataset of protein complexes upon which AlphaFold 3 and AF3Complex were evaluated was constructed using the Protein Data Bank (PDB) [14]. Initially, following the framework outlined in the AlphaFold-Multimer paper, proteins in the PDB were filtered based on the following criteria: Filtered proteins must have less than 3,000 deposited residues, have been released after September 30th, 2021, have at least two substituent protein chains, have at most nine substituent protein chains, not be monomeric, and have an experimentally determined template structure [7]. These filters ensured that the protein complexes were within the processing capabilities and specifications of AlphaFold 3 and AF3Complex. They also ensured that the proteins were not included within the AlphaFold 3 training datasets, for which the protein-inclusion cutoff date was September 30th, 2021 [8].

Next, after the initial filtering, using the *MMseqs2* package, the dataset of protein complexes was grouped into clusters based on a minimum sequence identity of 40%, as in the AlphaFold-Multimer paper, which ensures that the dataset is not saturated with similar protein motifs, thereby making each protein the models are evaluated upon sufficiently distinctive [7, 15]. From these clusters, a single representative was randomly chosen.

Lastly, to avoid trivializing the test by including protein structures that any of the models may have seen before, a similarity search between the low-homology protein complexes and all proteins in the PDB was undertaken. If any low-homology protein complex shared at least a 40% sequence similarity with a given protein in the PDB released on or before the cutoff date of September 30th, 2021, and could thus be plausibly in the AlphaFold 3 training dataset, it was removed. This last filtering step yielded a final dataset of 1091 protein complexes dissimilar to those on which AlphaFold 3 was trained and properly vetted for use in the forthcoming analysis.

For the protein complexes curated for this dataset, if the PDB template structure of the complex contained bound ligands and ions, they too were included in the dataset in the quantities found in the structure.

### Human Peptide Dataset Curation

To gauge the capacity of AF3Complex to model the interactions between proteins and peptides, which are often challenging for such models due to their diminutive size, a dataset of 91 human protein-peptide complexes was assembled from the PDB. Like the previous protein complex dataset, all of these complexes fell within the parameters of the guidelines outlined in the AlphaFold-Multimer paper, which are listed above, and were released after the AlphaFold 3 training dataset cutoff date of September 30th, 2021 [7, 8].

If the PDB template structure of the complex contained bound ligands and ions, they too were included in the dataset in the quantities shown in the structure.

### Curation of Antibody-Antigen Datasets

To assess AF3Complex’s capability to model the interactions between antibodies and antigens, two datasets from previous studies examining antibody-antigen modeling were collected. The first dataset was from a study examining AF2Complex’s performance on this protein complex variety and is a dataset of 36 different antibodies interacting with the SARS-CoV-2 spike RBD [12]. The second dataset is a collation of 108 various bound antibody-antigen complexes taken from a previous study on AlphaFold 3’s ability to predict antibody docking [16]. These datasets, in their respective studies of origin, were curated so that there would be no significant overlap between the complexes within them and those AlphaFold 3 was trained on, thereby avoiding trivialization of the task [16]. The first and second datasets will be referred to as the Gao and Hitawala datasets, respectively, hereafter.

### CASP16 Released Protein Complex Dataset

With the exclusion of two protein complexes that were too large for AF3Complex to process on the computing resources available, the dataset of nine protein complexes assembled from CASP16 were chosen as they were the only proteins with publicly available experimental structures to enable model assessment. The proteins in this dataset were T1201, H1202, H1204, T1206, T1234, T1235, H1232, H1233, and H1236.

When AF3Complex was employed to predict the structures of these protein complexes, it only had the information available to the participants in CASP16: the sequence and the stoichiometry of the protein complex. No other information was given to the model. Moreover, for the purpose of ranking the structures generated by AF3Complex, the phase of competition with the models achieving the highest scores was used for comparison.

### AF3Complex Predictive Metrics

A previous study introduced the Interface Similarity Score (IS-score) metric to more accurately and sensitively measure the structural similarity between protein complex interfaces [17] This score was developed to ameliorate the shortcomings associated with the TM-Score, which did not properly focus on the structural similarity of interfacial residues [9, 17]. In that study, a modified version of the IS-score was employed to function as a predictive metric within the model, thereby facilitating the model’s ability to choose the best structure from those generated.

We define the pIS by first introducing an intermediate metric, the predicted interface TM-Score (piTM), which is defined as follows:

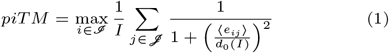

where ℐ is the set of interfacial residues within the predicted protein complex structure, and the cardinality of ℐ is the total number of residues *I* ≡ |ℐ| [4] Using the AlphaFold 3 confidence head, we can arrive at an estimate for the distance ⟨*e*_*ij*_ ⟩ between the interfacial residue and its assumed position in the experimental structure [8] The piTM score optimizes the position of the proteins in the complex, and *d*_0_(*I*) is the normalization factor that is defined as:

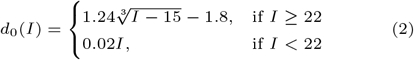

We derive the pIS from the piTM as follows:

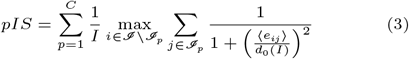

where we calculate the piTM for each protein chain *p* within the complex independently, adding together the values. In each chain, *p*, there are a certain number of interfacial residues, ℐ_*p*_, and ℐ represents the union of ℐ_*p*_.

The major difference between the pIS metric and the ipTM metric, the default metric for protein complexes in AlphaFold 3, is that the former focuses solely on interfacial residues rather than entire chains, which is preferable when trying to improve protein complex structure prediction [4, 7, 8].

Other, more recent, studies have offered possible alternatives to the AlphaFold 3 scoring metrics as well [18, 19]. Though these metrics have commonalities with the IS-score, their approach differs fundamentally in their sensitivity to different features of a protein complex. The pIS score, we believe, focuses more on the relevant regions of the complex for proper complex prediction, though such a claim would need to be further validated empirically.

Note that in the AF3Complex pipeline, the pIS metric is used to rank the generated structures to choose the best one, but this ranking metric is augmented by an auxiliary clash penalty, the same as the one used by AlphaFold 3, to ensure that the model never outputs structures with overlapping chains or residues [8]. Additionally, AlphaFold 3 does not use the ipTM score alone as its native ranking, rather, it uses a linear combination of the ipTM and pTM scores, as well as the aforementioned clash penalty and a supplementary reward for representing low-confidence regions in the generated structure as disordered regions [8].

### Model Testing Procedure

For each protein in the protein complex dataset, AlphaFold 3 and AF3Complex were provided with the same model seed, thereby keeping the internal workings of both models constant and allowing variation to stem only from the modifications made to the MSA pairing algorithm, the prediction metric calculation, and the ligand and ion data processing [8]. Both models were also given the same number of recycles, ten, and the same number of structural models to generate, five.

Similarly, when the AlphaFold 3 and AF3Complex models were fed data with ligands and ions purposefully excluded to examine to what degree such data impacts the accuracy of either model, the same MSAs and templates were employed as used in the inference pipeline for the models with ligand and ion data. This is permissible since ligand information is separate from the MSAs and template structures for a given protein [8].

### Statistical Tests

To evaluate the validity of the hypothesis that the model quality is improved using a certain scoring metric–the DockQ score, for example–we utilized a Wilcoxon signed-rank test, a non-parametric statistical test for distributions that do not follow a normal distribution, as is the case with the protein complexes [4]. Each of the tests was paired, as all the models make predictions on the dataset of protein complexes, and each of the tests was one-tailed.

## Results

### Model Comparison on Protein Complex Dataset

We first tested the performance of the AlphaFold 3 and AF3Complex models on the 1091 multimeric proteins within the novel protein complex dataset assembled in this study. For each protein complex, the models were given the same input–the amino or nucleic acid sequence and the ligand or ion information, if there was any–and were tasked with producing the best structural prediction possible.

Overall, the interfacial structural predictions of the AF3Complex model outperformed those of the AlphaFold 3 model, with a mean and median DockQ of 0.586 and 0.722 for AF3Complex and 0.573 and 0.704 for AlphaFold 3. A paired one-tailed Wilcoxon’s signed rank test on the DockQ distributions yielded a P-value of 4.2 × 10^−3^, indicating a statistically significant difference in the performance of the two models in terms of interfacial structural accuracy. The relative performance of the AF3Complex model relative to AlphaFold 3 based on DockQ can be viewed in Figure 2A and Supplementary Figure S1A.

**Fig. 2.**
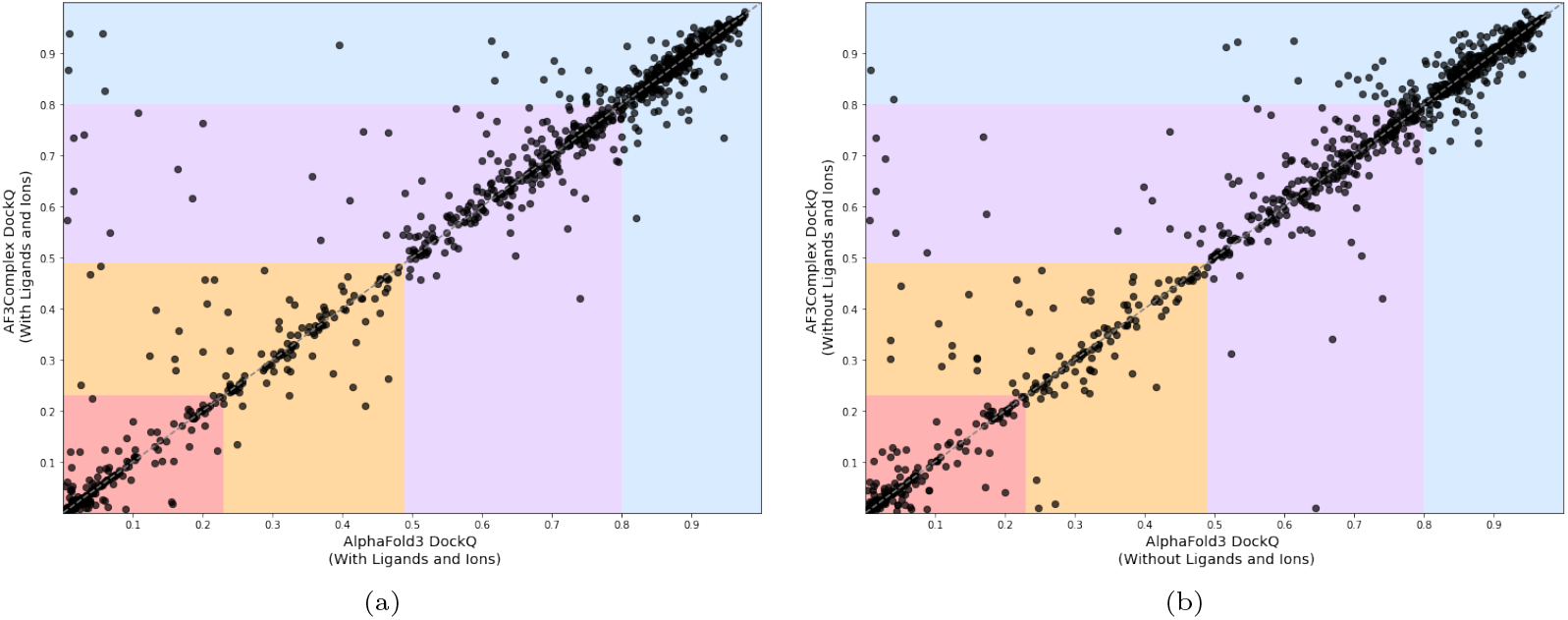
The figures show the relative DockQ scores of variations of AF3Complex and AlphaFold 3, where each dot is a single protein complex and the x-axis is the AlphaFold 3 model and the y-axis is the AF3Complex model. Each colored region represents the DockQ score distributions: Red [0.00, 0.23) is the incorrect region, Orange [0.23, 0.49) is acceptable accuracy, Purple [0.49, 0.80) is medium accuracy, and Blue [0.80, 1.00] is high accuracy. Figure (a) shows the performance when both models use ligand and ion data. Figure (b) shows the performance without this data.

When excluding all complexes for which both AF3Complex and AlphaFold 3 generated improper structures—defined as those with a DockQ value below 0.23—thus removing complexes that the AlphaFold 3 diffusion-model backbone struggles to model accurately, a statistically significant difference in TM-scores is observed between the two models, with a Wilcoxon signed-rank test yielding a P-value of 4.6 × 10^−3^. Specifically, AF3Complex achieves a mean and median TM-score of 0.898 and 0.970, respectively, and AlphaFold 3 achieves mean and median values of 0.888 and 0.968, respectively. This difference complements an increased performance gap in DockQ scores, as AF3Complex attains mean and median DockQ values of 0.735 and 0.796, respectively, while AlphaFold 3 reaches mean and median values of 0.718 and 0.790, respectively.

### Ablation Study With Ligand and Ion Data

In the previous analysis, both models utilized information about the number and types of ligands or ions in the protein complex they were tasked with modeling, as provided in the PDB. In reality, one often does not know in advance the quantities and varieties of ligands or ions bound to a protein complex, thus making the gauging of the model without this additional data valuable to understand their performance in practical application [5]. Hence, both models were reevaluated on the 1091-constituent protein complex dataset but, this time, with all ligand and ion data excluded from inputs.

The results of this evaluation showed the same trend as the previous: AF3Complex outperformed AlphaFold 3 in terms of interfacial structural accuracy, achieving a mean DockQ of 0.575 and a median of 0.702, while the latter model scored a mean DockQ of 0.563 and a median of 0.686. The distributions of scores achieved a P-value of 4.7 × 10^−5^ according to the same one-tailed Wilcoxon’s signed rank test as conducted earlier in the study, demonstrating that AF3Complex outperforms AlphaFold 3 even without the added benefit of ligand information. The relative performance of each model according to DockQ in this analysis can be viewed in Figure 2B and Supplementary Figure S1B.

The AF3Complex model with ligand and ion information does, however, eclipse both these models in performance, as can be seen in Supplementary Figure S2A, with a P-value of 1.3 × 10^−2^ when compared to its counterpart with ligand and ion data excluded, emphasizing the improvements yielded by the algorithm tasked with discerning whether ligand and ion information degrades the integrity of the predicted structural model.

A comparison of AlphaFold 3 models with and without ligand and ion information also reveals a clear performance boost in interfacial structure prediction when this data is included. The enhanced model outperforms the one without ligand and ion data, as demonstrated by a Wilcoxon signed-rank test with a P-value of 5.6 × 10^−4^ for DockQ scores. However, the extent of this improvement, reflected in mean and median DockQ scores of 0.573 and 0.704 with ligand and ion data, compared to 0.563 and 0.686 without, is less pronounced than the performance gap observed in the corresponding AF3Complex analysis. Supplementary Figure S2B shows the DockQ score distributions for both AlphaFold 3 model variants. This pattern suggests a possible broader trend among AlphaFold 3-based models, in which the presence of ligand and ion information generally enhances the accuracy of predicted protein complex structures. One possible explanation is that ligands and ions act as structural scaffolds that help guide protein conformation, thereby improving modeling. Alternatively, the model’s training on ligand and ion data may predispose it to perform better when such information is available.

### Analyzing Purposeful Ligand and Ion Data Omission in AF3Complex

Having shown that ligand and ion data improve AF3Complex’s performance, a natural question arises: why does the model exclude this critical data from some protein complex inputs based on relative pIS? The original intuition was that such data might add noise—forcing the model to handle not just protein and nucleic acid structure, but also ligand docking—-potentially degrading structural quality.

To assess this assumption, we examined the 72 protein complexes from the 1091-member dataset where AF3Complex excluded ligand and ion data due to a lower pIS. See Supplementary Figure S3 for the relative pIS scores. Of these 72 structures, 40 were better without the ligand data—indicating the model made the correct choice nearly 60% of the time. On average, these 72 complexes saw a DockQ improvement of 0.03, constituting a meaningful gain in accuracy.

Setting a threshold of 0.01 for absolute DockQ difference to identify consequential structural changes, 43 structures met this bar. Of the 43, 28 improved without ligand and ion data, and 15 improved with it, with an average DockQ gain of approximately 0.06. Raising the threshold to 0.05, 18 models qualified–14 of which outperformed their ligand-including versions, with an average DockQ improvement of approximately 0.13.

These results support AF3Complex’s comparative strategy for deciding when to include ligand and ion data.

Notably, most of the 72 complexes that excluded ligand and ion data were small with only two chains and fewer than 1000 residues. We speculate that in smaller models, ligand effects are minimal and add unnecessary complexity. In contrast, larger complexes benefit more from ligand inclusion, as ligands help reduce error propagation across larger surface areas and improve interfacial conformation.

### AF3Complex Evaluation on Human Peptide Dataset

A complicated yet vital form of protein complex prediction is the situation when a small sequence of amino acids, a peptide, interacts with one or more larger proteins [20]

AF3Complex was employed to predict the structure of 91 different human protein-peptide complexes to assess its ability to model this non-standard protein complex. The distribution of the DockQ values for AF3Complex and AlphaFold 3 on these 91 structural models can be seen in Figure 3A. When compared to the protein complex templates for the dataset, the structural predictions generated by the AF3Complex proved to be highly accurate, with a mean and median DockQ score of 0.642 and 0.747 across all models, indicating that the models were similar to their native structure in terms of their interfacial structures. Moreover, of the 91 structures generated, only 7 of the models were incorrect according to the DockQ value. More than two-thirds of the models were of medium quality or higher, with 38 models of high quality and 28 models of medium quality [11]. AF3Complex also outperformed AlphaFold 3 on this dataset, with the latter model achieving mean DockQ values of 0.626 and 0.697 on the dataset.

**Fig. 3.**
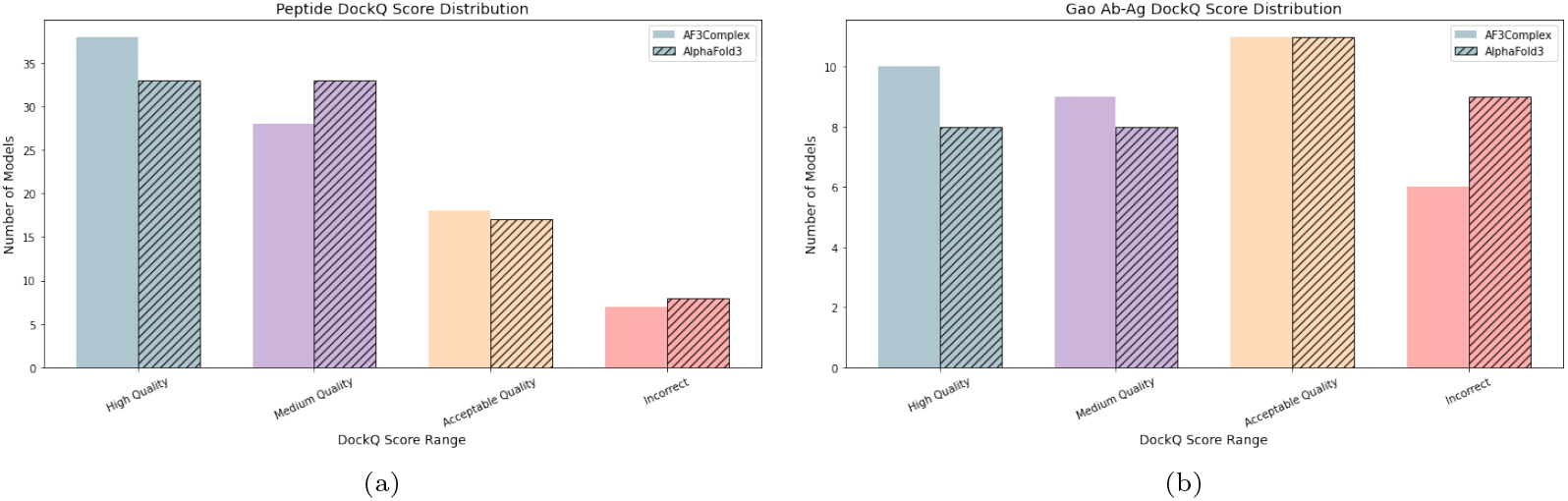
The figures display the number of generated structures that fall into the four aforementioned DockQ score regions for the 91-constituent protein-peptide complex dataset and the 36-constituent Gao antibody-antigen dataset. The solid colored bars depict AF3Complex’s performance, while the striped bars show AlphaFold 3’s performance. Figure (a) displays the quantities for the protein-peptide dataset, while figure (b) displays the quantities for the Gao antibody-antigen dataset.

**Fig. 4.**
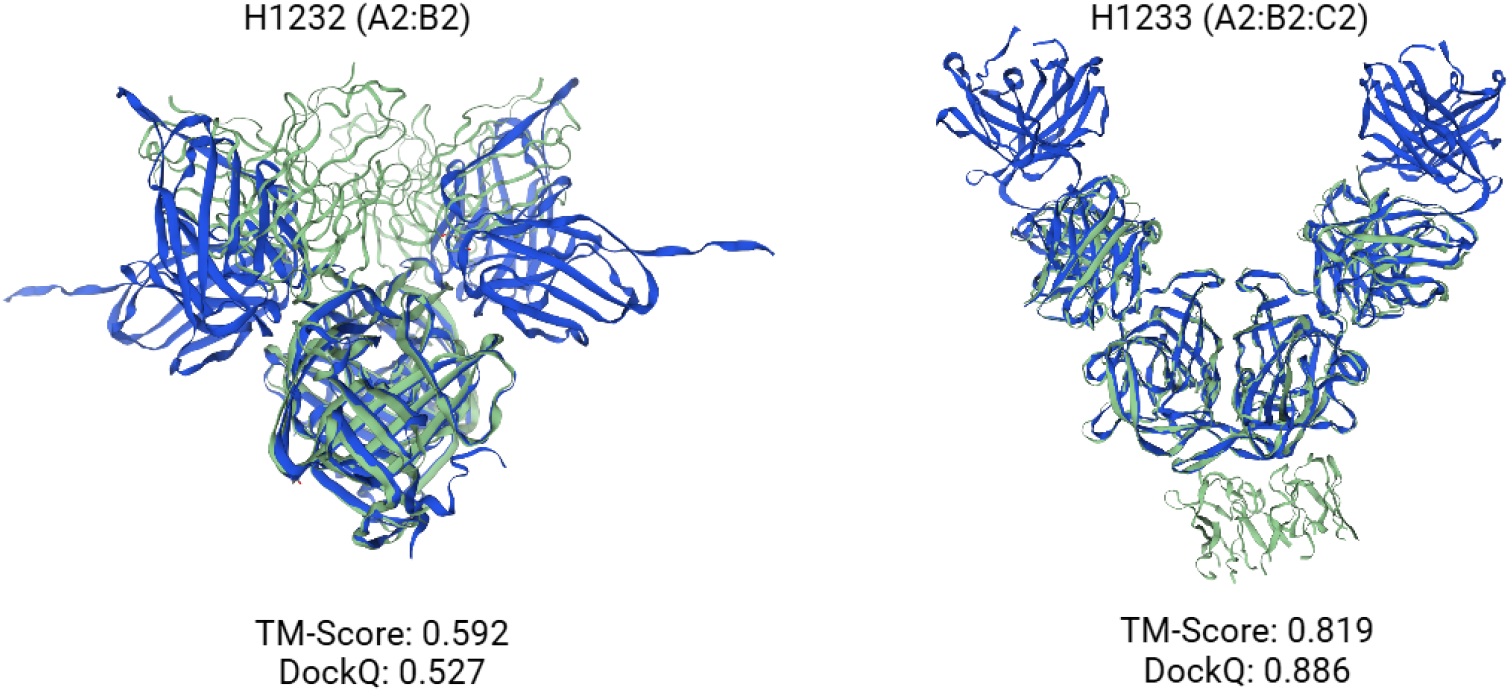
The structural predictions generated by AF3Complex on the two antibody-antigen complexes from CASP16 overlayed on their native structures with the associated TM-Score and DockQ metric results. The models generated by AF3Complex are colored blue while the native models are green.

Overall, the AF3Complex’s performance on the protein-peptide complex dataset shows a prodigious understanding of the structures.

### AF3Complex Evaluation on Antibody-Antigen Datasets

The prediction of antibody-antigen complex structures, which has numerous practical biomedical applications, is a well-known challenge for structural prediction models [21] Two datasets from previous studies benchmarking AF2Complex and AlphaFold 3 on antigen-antibody complex prediction were used as benchmarks [12, 16]. Using AF3Complex, new structural models were generated for the 36 complexes from the Gao dataset and the 108 complexes from the Hitawala dataset. The distribution of the DockQ values for the structural models generated by AF3Complex and AlphaFold 3 on the Gao dataset can be seen in Figure 3B, and the distribution for the Hitawala dataset can be seen in Supplementary Figure S4.

On both datasets, the structures generated by AF3Complex proved to be quite accurate when compared to their respective template structures, scoring a mean DockQ of 0.537 on the Gao dataset and 0.441 on the Hitawala dataset. Though the DockQ scores are not as high as those achieved by the model for the previous cases of general protein complexes and protein-peptide complexes, they represent an improvement over AlphaFold 3, which, on the very same datasets, achieved a mean of 0.492 and 0.424, respectively.

### AF3Complex Evaluation on CASP16 Protein Complexes

Following AF3Complex’s impressive performance when modeling the protein-peptide and antibody-antigen complexes, it seemed appropriate to examine how AF3Complex would perform relative to other structural prediction models on the protein complexes from the recent CASP16 competition. Hence, AF3Complex was used to model nine protein complexes from the CASP16 competition with publicly available template structures.

Overall, AF3Complex performed very well when predicting the structure of these protein complexes, generating two protein structures that proved better than any structures generated in the competition without human intervention, according to the weighted average of the DockQ metric. AF3Complex’s structures were consistently ranked within the top eight structures generated without human intervention by weighted average DockQ, except for one prediction on the T1235 protein complex, a highly challenging protein target, that was ranked eleventh. The structural predictions of AF3Complex on the two antibody-antigen complexes in the dataset-structures that would have placed second amongst the models generated without human intervention-can be seen in 4.

When averaged, the individual rankings by weighted average DockQ for each of the protein complex structures generated by AF3Complex, including those of the antibody-antigen complexes, yield a mean ranking of approximately 4.2. Comparing this to the average ranking of the best model for multimers in the competition not reliant on human intervention, Yang-Multimer [22], which had an approximate average ranking of 6.9 on these protein complexes, one can see that AF3Complex provides a sizeable increase in accuracy relative to the models used in CASP16.

## Discussion

The findings of this study clearly demonstrate that the downstream modification of AlphaFold 3 can lead to improvement in the model’s protein complex modeling capabilities. Two of the three modifications, the omission of the paired MSA and the use of the pIS metric, were based on modifications previously made to AlphaFold 2 to create AF2Complex [4]. The final modification, however, was novel to this study and took advantage of the AlphaFold 3 model’s unique ability to model ligands and ions by providing AF3Complex with two different inputs: one with the ligand and ion information and one without [8]. This approach allows the model to determine, through pIS comparison, whether ligand or ion information improves the quality of the structural model or detracts from it, thereby minimizing irregularities in the interfacial structures.

Through rigorous benchmarking of AF3Complex and AlphaFold 3 on different types of protein complexes, it is demonstrated that AF3Complex is at the state-of-the-art for protein complex prediction.

All three modifications contribute to the increase in performance we see in AF3Complex relative to AlphaFold 3. The modified MSA generation algorithm, as a previous study employing AF2Complex showed, omits paired MSAs, which allows the model more freedom to explore the structural and energetic relationships between amino acids in a protein structure and avoids many of the pitfalls of examining evolutionarily orthologous sequences [4]. It is important to note, however, that the AlphaFold 3 model architecture, which underpins AF3Complex, employs the information provided by MSAs to a much smaller degree than AlphaFold 2, on which AF2Complex is based [4, 8]. This architectural change reduced the impact of excluding paired MSAs.

Similarly, the revised predictive metric used in AF3Complex, pIS, was shown to improve the accuracy of structural predictions in AF2Complex relative to AlphaFold-Multimer [4]. A similar result can be seen in the analysis comparing the accuracy of the ranking metric employed by AlphaFold 3 and AF3Complex’s metric, pIS, described in the Supplementary Information, which showed that pIS more frequently chooses better structures from among those generated by AF3Complex. Ultimately, the pIS metric is constrained by the predicted aligned error matrices output by the internal AlphaFold 3 model’s confidence head, which is dictated by the model’s training, but, despite this, measurably improves the accuracy of the model’s predictions [4, 8].

Lastly, through an ablation study, we saw that the inclusion of ligand and ion data improved the accuracy of the AF3Complex model, allowing it to outperform AlphaFold 3. Further, even without the ligand and ion data, the other modifications made to AF3Complex allow it to perform just as well as AlphaFold 3 with that data given to it. Furthermore, we found that when AF3Complex omits ligand and ion data from the model inputs based on its predictive metrics, it more often yields a better structure for the protein complex. Omission of ligand and ion data was largely limited to smaller proteins of fewer than a thousand residues, for which we speculate the inclusion of ligand and ion data does less to stabilize the overall structure as compared to larger multimeric proteins—where small errors can propagate over a large surface area.

We also observed that AF3Complex also performs well for other cases besides the modeling of multimeric protein complexes. On a dataset of 91 protein–peptide complexes, the model achieved high interfacial accuracy. Additionally, across two separate antibody–antigen datasets [12, 16] AF3Complex demonstrated consistently strong performance, despite the notorious diversity and complexity of these interactions. Notably, in both the protein–peptide and antibody–antigen cases, AF3Complex outperformed AlphaFold 3. However, AF3Complex performed much better on the peptide dataset than on the antibody-antigen datasets, which is to be expected, as the former type of complex is generally less challenging than the latter.

Ultimately, AF3Complex is a derivative of AlphaFold 3 that builds upon that model’s ability to model protein complexes with nucleic acids, ligands, and ions. Often, but not always, it can provide highly accurate protein complex structures that exceed those of the model from which it was derived. This tool should have widespread applications in both research and clinical science.

## Competing interests

No competing interest is declared.

## Author contributions statement

J.F. and J.S. designed the research, J.F. wrote the source code, J.F. performed research and analyzed the data, J.F. prepared the first draft of the manuscript, J.F. and J.S. revised and proofread the manuscript.

## Acknowledgment

This research was supported in part by a grant GM 118039 from the Division of General Medical Sciences of the National Institutes of Health. A gift from the Ovarian Cancer Institute is gratefully acknowledged. The authors thank Google DeepMind for their work on AlphaFold 3. The authors would also like to thank Dr. Mu Gao, Davi Nakajima An, and Dr. Jerry M. Parks for their work developing AF2Complex.

## Notes

### Competing Interest Statement

The authors have declared no competing interest.

### Summary of Updates

The manuscript has been updated to reflect results on the new, expanded dataset and to enhance the manuscript's overall flow.

