## Supplementary Information for "AF3Complex Yields Improved Structural Predictions of Protein Complexes"

### 1 Examination of the pIS Metric

In addition to understanding how the exclusion of ligand and ion data affects the accuracy of the AF3Complex model, we also wanted to examine to what degree the pIS scoring metric improves the accuracy of the structural models. Though this scoring metric was previously shown to improve the outputs of AF2Complex [1], since the AlphaFold 3 model from which the predicted aligned error matrices are derived is different architecturally and in terms of training [2], it is necessary to examine the impacts of this metric specifically for the AF3Complex model, which is reliant upon AlphaFold 3.

For each protein complex in the 1091-constituent protein complex dataset, we examined whether the best model chosen by the ranking score metric used in AlphaFold 3, which is a linear combination of ipTM and pTM values [2], and the metric employed by AF3Complex, pIS, correctly reflected the best model based on the DockQ score. Of the 1091 protein complexes, we found that the pIS had a marginally better performance, choosing the best model 287 times on the dataset with ligand and ion information included, while the AlphaFold 3 ranking score did so 283 times. However, if we examine the number of times the metrics chose a better model according to DockQ relative to each other, regardless of whether it was the best model overall, the pIS metric chose a better model 305 times, and the traditional ranking score did so only 256 times.

It is important to note that the improvement yielded by the modified predictive metric is fundamentally limited by the training of the model upon which it operates, and this explains the high rates of concordance across metrics. Though pIS better reflects the protein complex structures it aims to predict, like the AlphaFold 3 ranking score, is still wholly reliant on the outputs of the AlphaFold 3 confidence head, which cannot be modified without changing the model's weights.

### 2 Further Examination of 1091-Constituent Benchmark Results

Having established that AF3Complex outperforms AlphaFold 3 both with and without ligand information, we next sought to determine the conditions under which this improvement is most pronounced. As illustrated in Figures 2A and 2B and in Supplementary Figures S2A and S2B, the advantage conferred by AF3Complex is not universal. Applying a threshold of 0.05 or greater in DockQ improvement, we find that AF3Complex outperforms AlphaFold 3 on 140 complexes when ligand and ion information is included, and on 150 complexes when such information is omitted. The distribution of DockQ scores across CASP bins can be found in Supplementary Figures S5A and S5B.

For both distributions—those with and without ligand and ion information—the majority of complexes showing a performance difference consist of fewer than 1,000 residues. This reflects the underlying composition of the dataset, which is predominantly made up of proteins under 1,000 residues, consistent with the general size distribution found in the PDB. Similarly, most of the complexes are dimers in both cases; this is expected given that dimers are the most common oligomeric state in both the dataset and the PDB at large.

### 3 The Role of Ligands in the 1091-Constituent Benchmark Results

Among the 140 protein complexes where AF3Complex outperforms AlphaFold 3 in the ligand-containing dataset, only 67 include ligand and ion information. Within this subset, no single type of ligand or ion predominates. Furthermore, the spatial positioning of these molecules within or around the complex is variable, and no clear relationship emerges between their location and the accuracy of the predicted interfacial structure. Amongst those models that have ligand and ion data excluded, there is no apparent global conformational change with or without such ligand data.

It is highly likely that there are certain characteristics of these proteins that prime them to have their respective structural predictions improved by the modifications made to create AF3Complex, but these are likely not apparent to an observer, as they are dependent on the complex internal system of the model.

Importantly, the presence of ligand and ion information does not guarantee improved performance. In fact, AF3Complex does not consistently outperform AlphaFold 3 even among the 579 complexes in the dataset that include such molecules. As shown in Supplementary Figure S6, there are several cases where AlphaFold 3 achieves higher interfacial accuracy than AF3Complex despite the presence of ligands or ions, though only to a minor degree.

For illustrative examples of structure predictions where the performance gap between AF3Complex and AlphaFold 3 is most pronounced, refer to Supplementary Figures S7 and S8.

### 4 Further Examination of Antibody-Antigen Benchmark Results

Across both the Gao and Hitawala antibody–antigen datasets, AF3Complex consistently outperforms AlphaFold 3. The improvement is especially pronounced in the Gao dataset, though this may be partly attributed to its structural redundancy, as all complexes involve the SARS-CoV-2 RBD. Nevertheless, the magnitude of improvement observed on this dataset is noteworthy.

In contrast, the performance gains on the Hitawala dataset are more modest but remain consistent. Unlike the Gao dataset, where a number of complexes show substantial differences in accuracy, the Hitawala dataset exhibits fewer such extremes. This likely reflects the Hitawala dataset's greater diversity. Still, there are select cases within the Hitawala set where AF3Complex achieves marked improvements over AlphaFold 3—one such example is shown in Supplementary Figure S9.

There may be additional opportunities to improve antibody–antigen structure predictions for both models. As noted by the developers of AlphaFold 3, increasing the number of inference seeds—up to 1,000 in their case—can lead to dramatic gains in performance for such complexes [2]. However, this approach is extraordinarily computationally intensive and was not feasible within the scope of this study.

### 5 Assorted Figures

**Fig S1A**

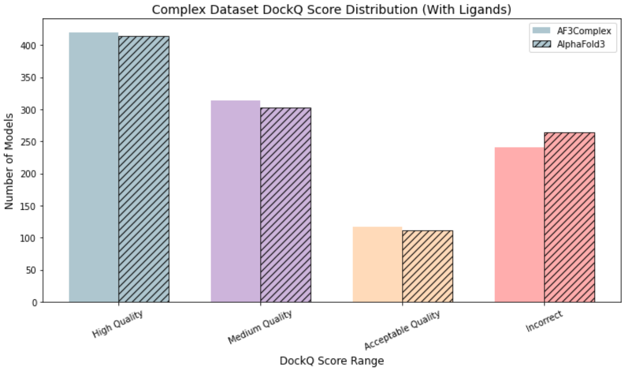


**Fig S1B**

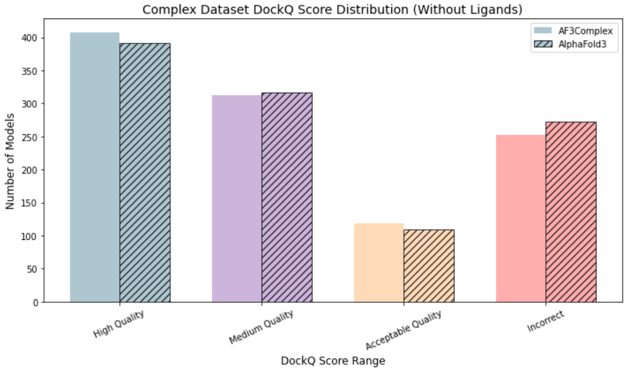


**Fig S1.** The figures display the number of generated structures that fall into the four DockQ score regions for the 1091-constituent protein complex dataset. The solid colored bars depict AF3Complex's performance, while the striped bars show AlphaFold 3's performance. Figure S1A displays the distribution for models that have access to ligand and ion information, while Figure S1B displays the distribution when they do not.

**Fig 2A**

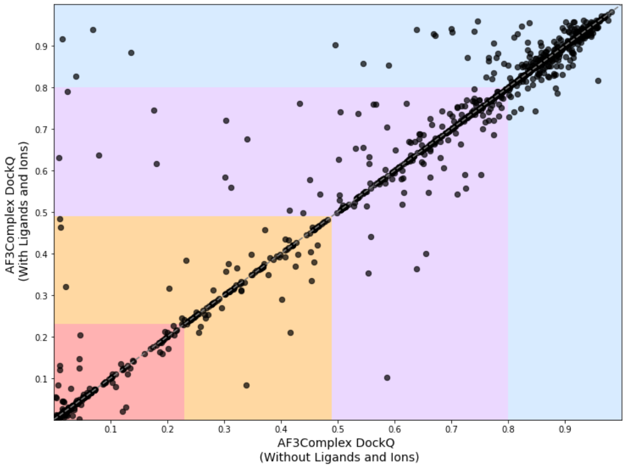

**Fig 2B**

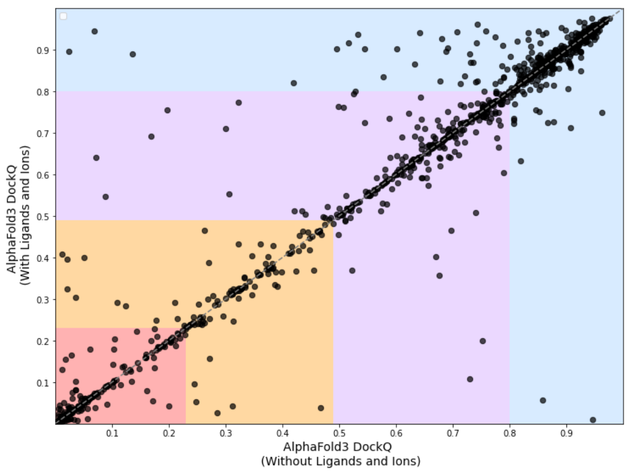


**Fig S2.** The relative DockQ scores of variations of AF3Complex and AlphaFold 3, where each dot is a single protein complex, the x-axis is the AlphaFold 3 model, and the y-axis is the AF3Complex model. Each colored region represents the DockQ score distributions: Red [0.00, 0.23) is the incorrect region, Orange [0.23, 0.49) has acceptable accuracy, Purple [0.49, 0.80) is medium accuracy, and Blue [0.80, 1.00] is high accuracy. Figure S2A shows the relative performance between AF3Complex with ligand and ion information and AF3Complex without that information. Similarly, Figure S2B shows the relative performance of AlphaFold 3 with ligand and ion information and AlphaFold 3 without it.

**Fig S3**

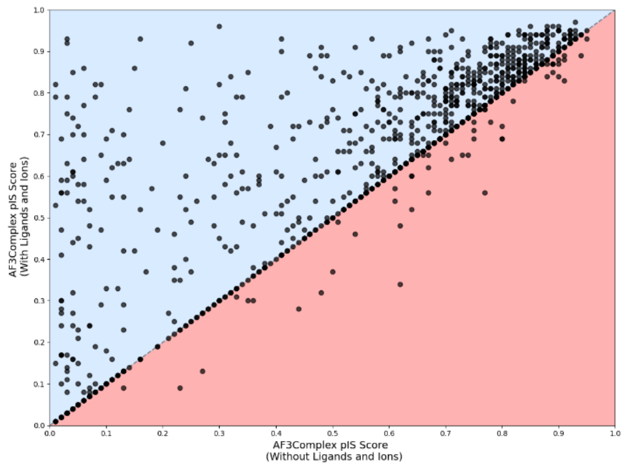


**Fig S3.** This figure shows the relative pIS scores of the AF3Complex outputs with and without ligand and ion information for each of the 579 protein complexes among the 1091-constitutent protein complex dataset, where each dot is a single protein complex, that have PDB-extracted ligand and ion information included in their input.

**Fig S4**

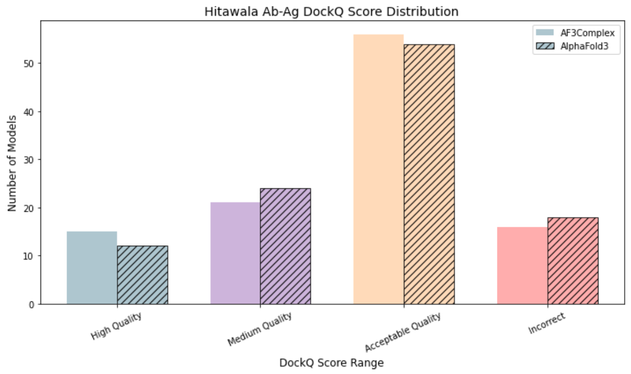


**Fig S4.** Number of generated structures that fall into the four DockQ score regions for the 108-constituent Hitawala antibody-antigen dataset. The solid-colored bars depict AF3Complex's performance, while the striped bars show AlphaFold 3's performance.

**Fig S5A**

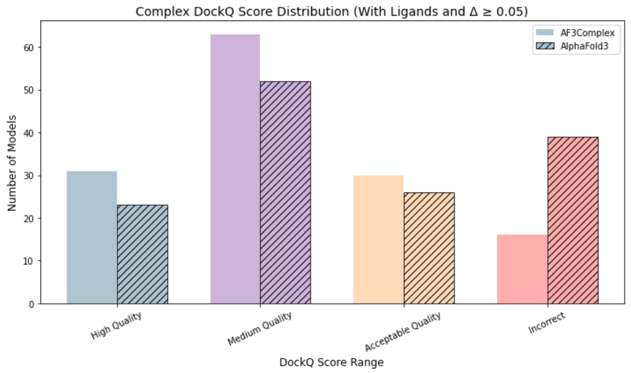


**Fig S5B**

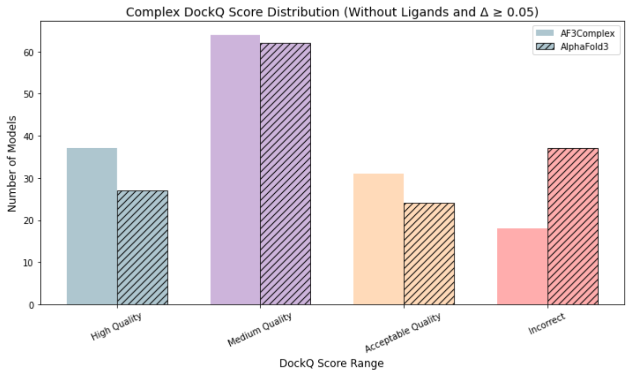

**Fig S5.** The number of generated structures that fall into the four DockQ score regions for the 1091-constituent protein complex dataset when a threshold of an absolute difference of at least 0.05 is applied. The solid-colored bars depict AF3Complex's performance, while the striped bars show AlphaFold 3's performance. Figure S5A displays the distribution when both models have access to ligand and ion information, while Figure S5B displays the distribution when they do not.

**Fig S6**

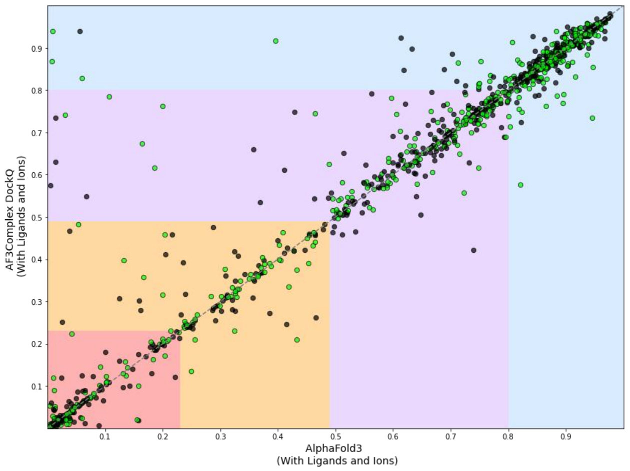


**Fig S6.** This figure shows the relative DockQ scores of variations of AF3Complex and AlphaFold 3, where each dot is a single protein complex, and the x-axis is the AlphaFold 3 model, and the y-axis is the AF3Complex model. Each colored region represents the DockQ score distributions: Red [0.00, 0.23) is the incorrect region, Orange [0.23, 0.49) is acceptable accuracy, Purple [0.49, 0.80) is medium accuracy, and Blue [0.80, 1.00] is high accuracy. In this figure, AF3Complex has ligand and ion information and so does AlphaFold 3.
The protein complexes that have ligand and ion information as part of their input features are highlighted in green. Those that do not are colored black.

**Fig S7**

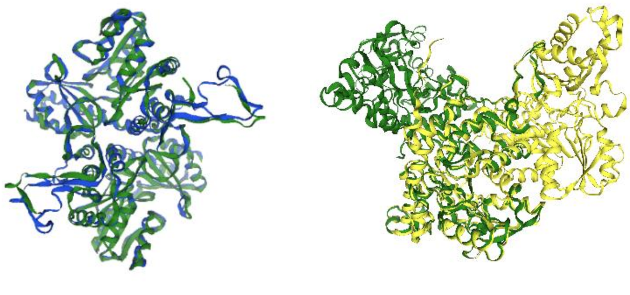

**Fig S7** These are the structural predictions generated for the protein complex 7YEO from the 1091-constituent protein complex dataset by AF3Complex and AlphaFold 3. The left structure is a superimposition of the structure generated by AF3Complex, colored in blue, and the native structure, colored in green. The right structure is a superimposition of the structure generated by AlphaFold 3, which is colored yellow, on the native structure, which is also colored green. This protein complex is the one on which AF3Complex most outperforms AlphaFold 3 in terms of interfacial accuracy when ligand and ion data is present, and it is visually evident that the structural alignment globally and interfacial similarity is much poorer for the AlphaFold 3 model.

**Fig S8**

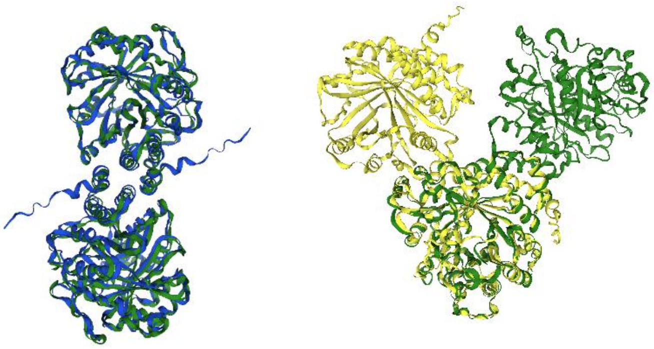

**Fig S8** These are the structural predictions generated for the protein complex 9IIA from the 1091-constituent protein complex dataset by AF3Complex and AlphaFold 3. The left structure is a superimposition of the structure generated by AF3Complex, colored in blue, and the native structure, colored in green. The right structure is a superimposition of the structure generated by AlphaFold 3, which is colored yellow, on the native structure, which is also colored in green. This protein complex is the one on which AF3Complex most outperforms AlphaFold 3 in terms of interfacial accuracy when ligand and ion data is not present, and it is visually evident that the structural alignment globally and interfacially is much poorer for the AlphaFold 3 model.

**Fig S9**

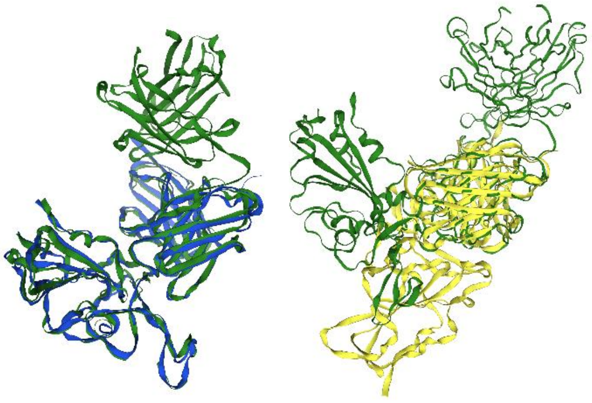

**Fig S9** This figure depicts the structural predictions generated for one antibody-antigen complexes from the 108-constituent Hitawala antigen-antibody protein complex dataset with PDB id 8GB6 by AF3Complex and AlphaFold 3. The left structure is a superimposition of the structure generated by AF3Complex, colored in blue, and the native structure, colored in green. The right structure is a superimposition of the structure generated by AlphaFold 3, which is colored yellow, on the native structure, which is also colored in green. This complex is the one on which AF3Complex most outperformed AlphaFold 3.
